# Identifying brain-penetrant small molecule modulators of human microglia using a cellular model of synaptic pruning

**DOI:** 10.1101/2024.12.23.630139

**Authors:** Liam T. McCrea, Rebecca E. Batorsky, Joshua J. Bowen, Hana Yeh, Jessica M. Thanos, Ting Fu, Roy H. Perlis, Steven D. Sheridan

## Abstract

Microglia dysregulation is implicated across a range of neurodevelopmental and neurodegenerative disorders, making their modulation a promising therapeutic target. Using PBMC-derived induced microglia-like cells (piMGLCs) in a scalable assay, we screened 489 CNS-penetrant compounds for modulation of microglial phagocytosis of human synaptosomes in a validated assay for microglia-mediated synaptic pruning. Compounds from the library that reduced phagocytosis by ≥2 standard deviations across the library without cytotoxicity were validated in secondary screens, with 28 of them further confirmed to reduce phagocytosis by 50% or more. Image-based morphological measurements were calculated to measure the degree of ramified vs. amoeboid morphotype as an indicator of activation state. Additionally, transcriptomic profiling indicated divergent effects on cell signaling, metabolism, activation, and actin dynamics across confirmed compounds. In particular, multiple CNS-penetrant small molecules with prior FDA approval or demonstration of safety *in vivo* demonstrated modulatory effects on microglia. These potential disease-modifying agents represent high-priority candidates for repositioning studies in neurodevelopmental, neuroinflammatory, or neurodegenerative disorders.

## Introduction

Microglia have essential functions in the developing and mature central nervous system (CNS).^1,2^ After populating the brain during early embryonic development^1,3^, they play a central role in immunosurveillance^4,5^, regulation of neuronal progenitor populations^6^, neurogenesis^7,8^, and shaping synaptic networks via selective pruning.^9-11^.

Convergent lines of evidence including animal studies, post-mortem and *in vitro* human models associate dysregulated microglia-mediated synaptic pruning with the pathophysiology of CNS disorders.^12-14^ In particular, recent investigation have shown that dysregulated microglial phagocytosis, such as during synaptic pruning, may be associated with neurodevelopmental disorders including bipolar disorder, autism and schizophrenia.^12,15,16^ Postmortem studies demonstrate reduced cortical dendritic spine density within the brains of individuals with schizophrenia^15-17^, consistent with clinical structural neuroimaging studies.^18,19^ Parallel lines of evidence from rodent models and human genomics suggest that these disorders may arise, in part, from dysfunctional microglia-mediated pruning.^14,20^ Microglial processes may amplify or exacerbate the downstream effects of these disorders even when such disease states are initiated by other processes.^21^

For example, microglia become activated upon initial accumulation of amyloid-beta (Aβ) plaques in Alzheimer’s Disease (AD), leading to an increased release of pro-inflammatory cytokines, reactive oxygen species (ROS), and other neurotoxic substances.^22^ This chronic neuroinflammatory response exacerbates neuronal damage and accelerates disease progression.^23^ Conversely, in a mouse AD model, inhibition of phagocytic microglia has been shown to reduce the extent of early synapse loss.^24^

Interventions that modulate microglial function are therefore promising candidates for disease-modifying treatments. While nonspecific anti-inflammatory agents have been investigated clinically with varied efficacy^22,25,26^, few small-molecule microglia modulators have been specifically identified, and even fewer have been investigated mechanistically.

*In vitro* assays utilizing human microglia represent a promising means for the discovery of new compounds or repurposing of existing therapeutics that modulate microglial function. However, due to their limited availability, microglia derived from post-mortem or surgical biopsy brain material lack the scalability to perform high-throughput screening (HTS). These limitations have led to development of alternative microglia-like cellular models, such as from the differentiation of human induced pluripotent stem cells (iPSCs).^27^ These iPSC-derived microglia models, however, also have limitations, including the extensive resources required for large scale *in vitro* derivation, expensive media over extended time in culture, and significant technical artifacts after scaling due to line-to-line variability. Another strategy utilizes the direct reprogramming of human peripheral mononuclear blood cells (PBMCs) to microglia-like cells (piMGLCs).^27-29^ The monocyte fraction of PBMCs possesses high phenotypic plasticity and can adopt a microglial-like phenotype *in vitro.* This direct conversion of human monocytes to induced microglia-like cells (piMGLCs) therefore represents a promising strategy for generating large-scale assays amenable to HTS from human blood.^14,28-30^

We scaled our high-content, image-based assay for functional synaptic pruning utilizing the engulfment of human neural culture-purified synaptic vesicles (synaptosomes) by piMGLCs ^14,30,31^ to screen a library of known CNS-penetrant compounds. To investigate mechanisms of action, we secondarily characterized morphology as well as altered transcriptomic pathways.

As many of these compounds have previously been investigated in human studies for other indications, with some already in clinical use, we aimed to identify high-priority molecules with known CNS penetrance and safety for repositioning in psychiatric and neurodevelopmental disorders.

## Results

### PBMC-derived induced microglia-like cells as a scalable HTS platform for identifying modulators of synaptosome phagocytosis

We have previously optimized and validated a cytokine induction method to derive PBMCs into piMGLCs that exhibit a high degree of transcriptomic similarity to primary microglia, immortalized primary microglia and isolated adult post-mortem microglia.^14,30^ For large-scale derivation of piMGLCs used in our assays, we generated a large biobank of assay-ready cells by isolating, aliquoting, and cryopreserving a single large batch of PBMCs (approx. 2 to 3x10^9^ cells) from a commercially available leukapheresis preparation (leukopak). This large batch size was used to minimize batch effects in the screening assays. From these banked PBMC aliquots, we used this cytokine induction method and scaled this process to 96-well format. piMGLC batches were induced for ten days before being harvested, pooled and replated into 96-well assay plates. After three days recovery in assay plates, the majority of piMGLCs displayed a ramified morphology (Fig. 1A), with positive immunostaining for the microglial markers IBA1, PU.1, CX3CR1 and P2RY12 (Fig. 1B). We have previously demonstrated that these piMGLCs demonstrate a high degree of transcriptomic similarity to primary microglia, immortalized primary microglia, and isolated adult post-mortem microglia.^30^

**Figure 1.**
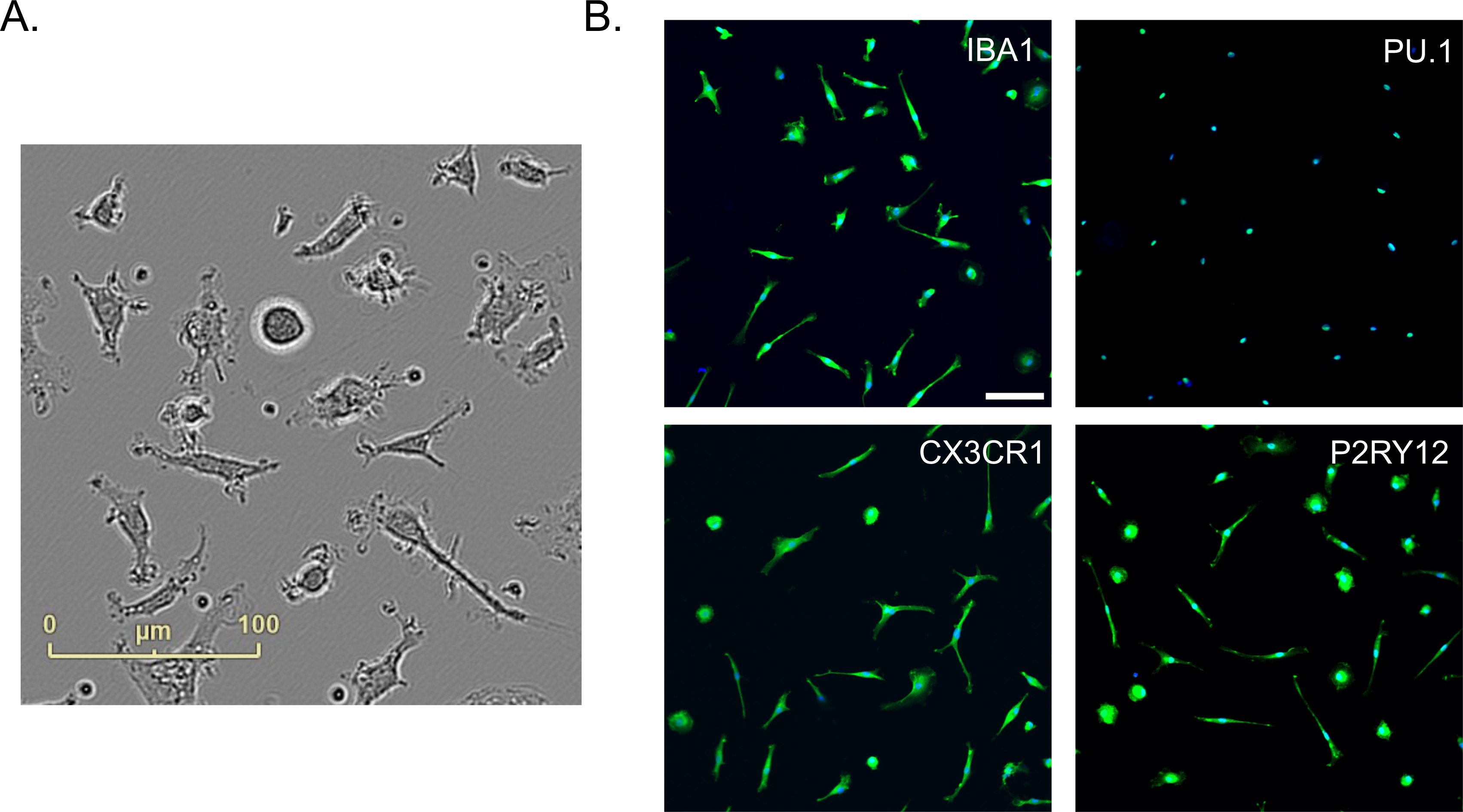
Derivation of PBMC microglia-like cells (piMGLCs). (a) Representative phase-contrast image of piMGLCs. Scale bar, 100 μm. (b) Confocal images of piMGLCs stained for indicated canonical microglial markers IBA1, PU.1, CX3CR1 and P2RY12. Scale bar, 100 μm

To screen for potential modulators of microglia-mediated synaptic pruning, we quantified phagocytosis of synaptosomes isolated from large-scale human iPSC-derived differentiated neuronal cultures and labeled with a pH-sensitive dye (pHrodo) that fluoresces in the acidic post-phagocytic phagolysosome compartments. We used these reagents in conjunction with the piMGLCs in an *in vitro* model of microglial synaptic engulfment (Fig. 2A). After incubation with synaptosomes for 3 hours, cells were fixed and stained with the microglial marker IBA1 to identify cellular perimeters. This cell segmentation, along with the quantification of bright red fluorescence within cells, indicating cellular uptake of pHrodo-labeled synaptosomes^14,30^, were performed using confocal microscopy images input into an automated image analysis pipeline in CellProfiler.^32^

**Figure 2.**
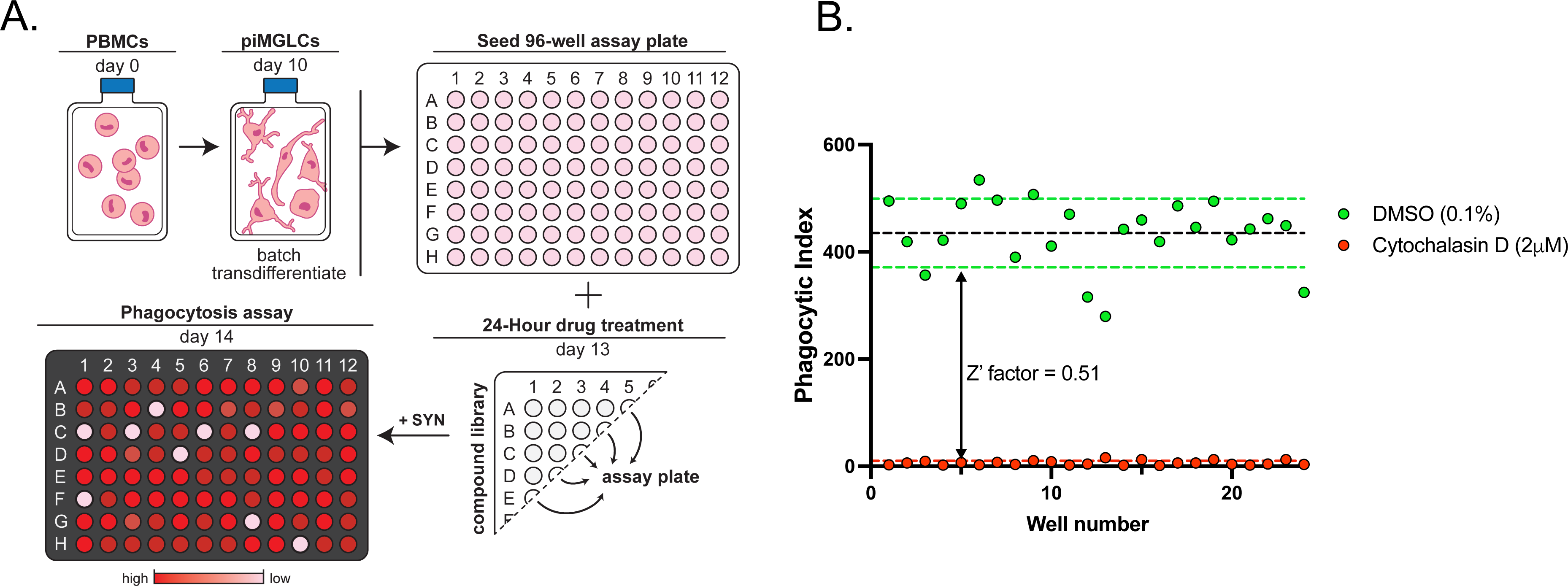
Image-based phagocytosis assay optimization for CNS-penetrant compound HTS screening. (a) Overview workflow for large-scale batch derivation of piMGLCs and 96-well functional synaptosome phagocytosis image-based screens. (b) Z’ factor = 0.51 was determined using 0.1% DMSO (green) as negative control and 2μM cytochalasin D (red) as positive control phagocytosis inhibitor, 24 wells of each. Data points represent phagocytic indices of each well, calculated as the sum synaptosome area divided by sum cell count within 12 image fields per well. Black dashed lines are the group means, and color dashed lines signify one standard deviation around the mean for the group of the corresponding color.

To determine assay robustness in 96-well plate format, we treated 24 wells each for 24 hours with either only 0.1% DMSO (negative carrier control) or 2uM cytochalasin D (CytoD), a cell-permeable inhibitor of actin polymerization, used as a positive control treatment to inhibit phagocytosis. Phagocytosis activity (expressed as phagocytic index, see Methods) in positive control and negative untreated wells was determined to have a robust Z-factor score of 0.51 (Fig. 2B), indicating a high-quality assay for screening. Z-factor represents a standard measure of statistical effect size in high throughput screening between compound ‘hits’ and background variation to determine assay robustness. ^33^

### Functional CNS-penetrant compound screening identifies compounds that modulate synaptosome phagocytosis

We performed image-based primary compound screening, allowing for the screening of compounds that affect microglia function as indicated by phagocytosis of isolated synaptosomes in a model of synaptic pruning.^14,30^ Using our optimized assay in 96-well plate format we screened a CNS-penetrant small-molecule library of 489 compounds (Supplemental Data Table S1) by treating at an initial concentration of 10uM for 24 hours before the addition of labeled synaptosomes to initiate the assay with a small number screened at 2uM due to their limited solubility in stocks (Supplemental Data Table S2). After 3 hours, cells were fixed and analyzed by confocal microscopy. Treated wells containing below a threshold number of remaining cells (<10% compared to DMSO) were considered to indicate possible toxicity. Other criteria for omission included two image quality control filters: one removing images with high cell densities that could not be accurately segmented, and another for extremely high synaptosome fluorescence due to segmentation of background signal or image artifacts. After excluding 35 compounds (7%) with these initial filters, 454 compounds remained for further phagocytic activity determination after eliminating these wells. We further prioritized compounds more stringently based on those that deviated from the mean phagocytic index of the DMSO negative control by at least a Z-score of +/-2, or two standard deviations (2SD), as initial primary hits. We did not identify any activators with a Z-score ≥ 2 using this method of prioritizing hits but were able to identify 47 compounds (∼10% of the entire library) that stringently (Z-score ≤ -2) inhibited phagocytosis of synaptosomes (Fig. 3A and Supplemental Data Table S3).

**Figure 3.**
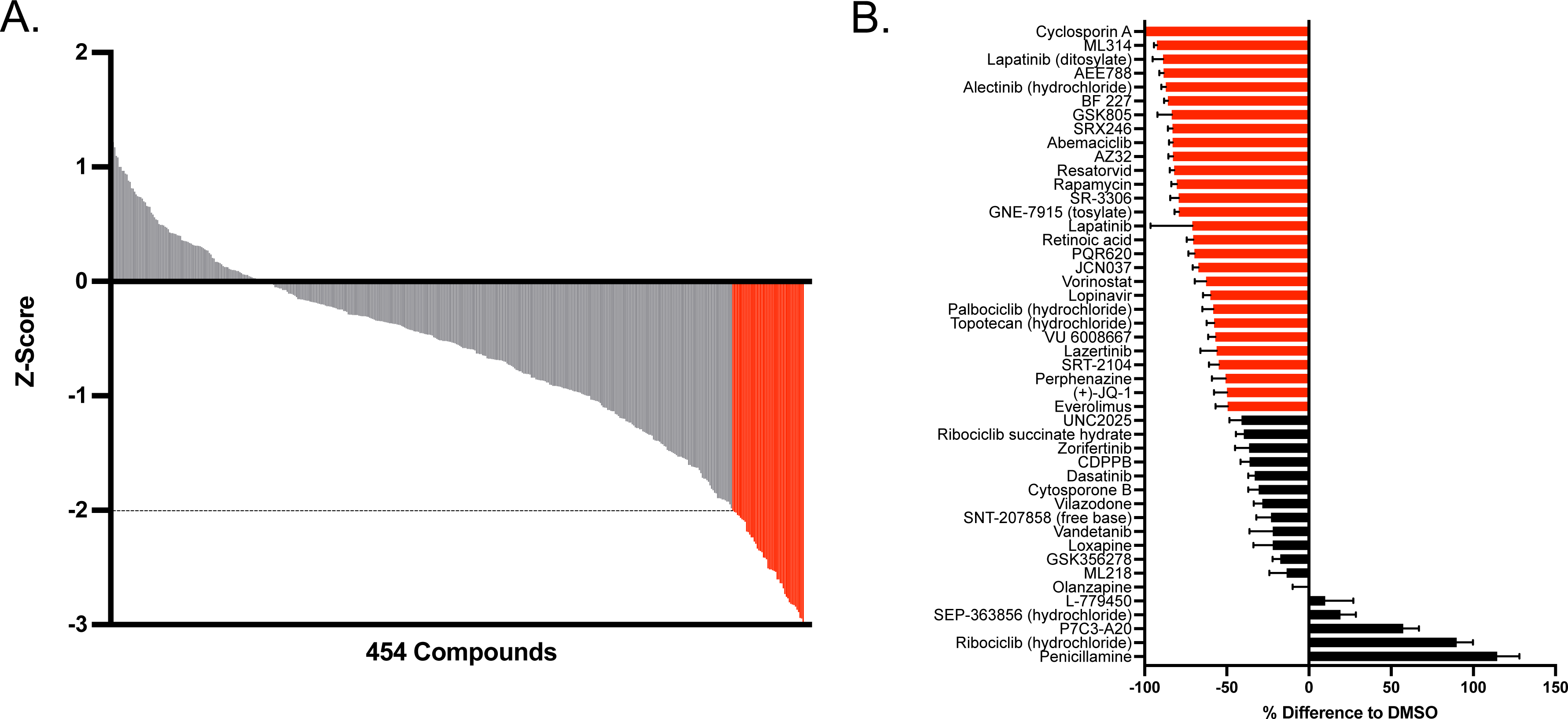
Image-based compound screen for synaptosome phagocytosis modulators in piMGLCs. (a) Primary screen of 489 CNS-penetrant compounds assayed at 10μM resulted in 454 measurable phagocytic activity (35 removed due to toxicity). 47 compounds that inhibited by a Z-score compared to DMSO (Z-score=0) of -2 or below indicated in red. (b) Secondary confirmation screen of hit compounds measured as the ratio of mean phagocytic index per image field (n=15) of indicated compound treatment to mean DMSO only control. Compounds with a ratio of 0.5 or less indicated in red. Error bars indicate SEM.

To confirm results from the primary library screen and further demonstrate independent reproducibility, we performed a secondary confirmation screen on the 47 hit compounds with freshly prepared bulk compound stocks. One compound, MK-28, was determined to be toxic, and six compounds were shown to either increase phagocytosis (penicillamine, ribociclib HCl, P7C3-A20, SEP-363856 and L-779450) or have no measurable effect (olanzapine) when compared to carrier (DMSO) negative control measures (Fig. 3B). While the remaining 40 compounds all reduced synaptosome phagocytosis as was observed in the primary screen, 28 compounds demonstrated a 50% reduction or greater when compared to the DMSO control confirming these compounds as inhibitors with apparent IC50 values of ≤ 10μM (Supplemental Data Table S4). The comparison of the measured phagocytic indices for the forty remaining inhibiting compounds between the primary and secondary screening results revealed a moderate correlation for most compounds (R=0.3, p=0.067 across compounds), with some outliers reflecting possible subtle differences between drug concentrations in initial primary screen compound library versus independent individual secondary screen compounds.

Many of the compounds confirmed to reduce microglial phagocytosis in our screen are current FDA-approved medications [14/28; 50%] or have been investigated in human studies [4/28, 14%] (Supplemental Data Table S5). Immunosuppressant compounds that show robust microglia phagocytosis reduction, such as cyclosporin A and rapamycin, are effective in preventing organ rejection in transplant recipients.^34^ Several identified kinase inhibitors in the screen are currently used for cancer treatment (e.g. lapatinib, alectinib, abemaciclib and recently FDA-approved lazertinib) either alone or in combination with other therapeutics. Perphenazine, loxapine and vilazodone have been indicated for psychiatric diagnoses including schizophrenia, bipolar disorder and major depression. While their primary mechanism of action is best understood for their effects on neurotransmission, substantially less is understood about the indirect effects of these compounds on microglial function. Conversely, 10/28 (36%) of confirmed hit compounds are investigational or experimental without any current clinical application. These include such the c-Jun N-terminal kinases (JNK) inhibitor SR-3306, investigated for dopaminergic neuron protection in animal models of Parkinson’s disease (PD)^35^ and the leucine-rich repeat kinase 2 (LRRK2) inhibitor GNE-7915, which along with its derivatives showed promising pharmacokinetic and safety profiles in rodent and non-human primate models for potential application of LRRK2 inhibition in PD.^36^

### Phenotypic compound screening identifies CNS-penetrant compounds that modulate microglial morphology

In the adult brain, microglia typically exist in a surveillant state characterized by a ramified morphology.^4^ Microglia undergo morphological changes into a more amoeboid shape upon activation due to various responses and disease states. This transformation can be used as a proxy for activation state ^37^, with more ramified or amoeboid morphologies representing a surveillant or more activated state, respectively.

An advantage of our image-based arrayed screen is its versatility, as it allows for the determination of multiple parameters of the cellular response in parallel to phagocytosis upon compound treatment. Using CellProfiler^32^, we examined cellular morphologic parameters in the IBA1+ stained cells used to quantify synaptosome uptake to additionally determine measures of solidity and eccentricity (see Supplemental Methods) in the compound-screened piMGLCs (Fig. 4). We have previously shown that these morphometric parameters allow for the high-throughput categorization of individual cells into ramified (low solidity, high eccentricity), amoeboid (high solidity, low eccentricity) and in-between bipolar, or rod-shaped (mid solidity, mid eccentricity) morphotypes.^31^

**Figure 4.**
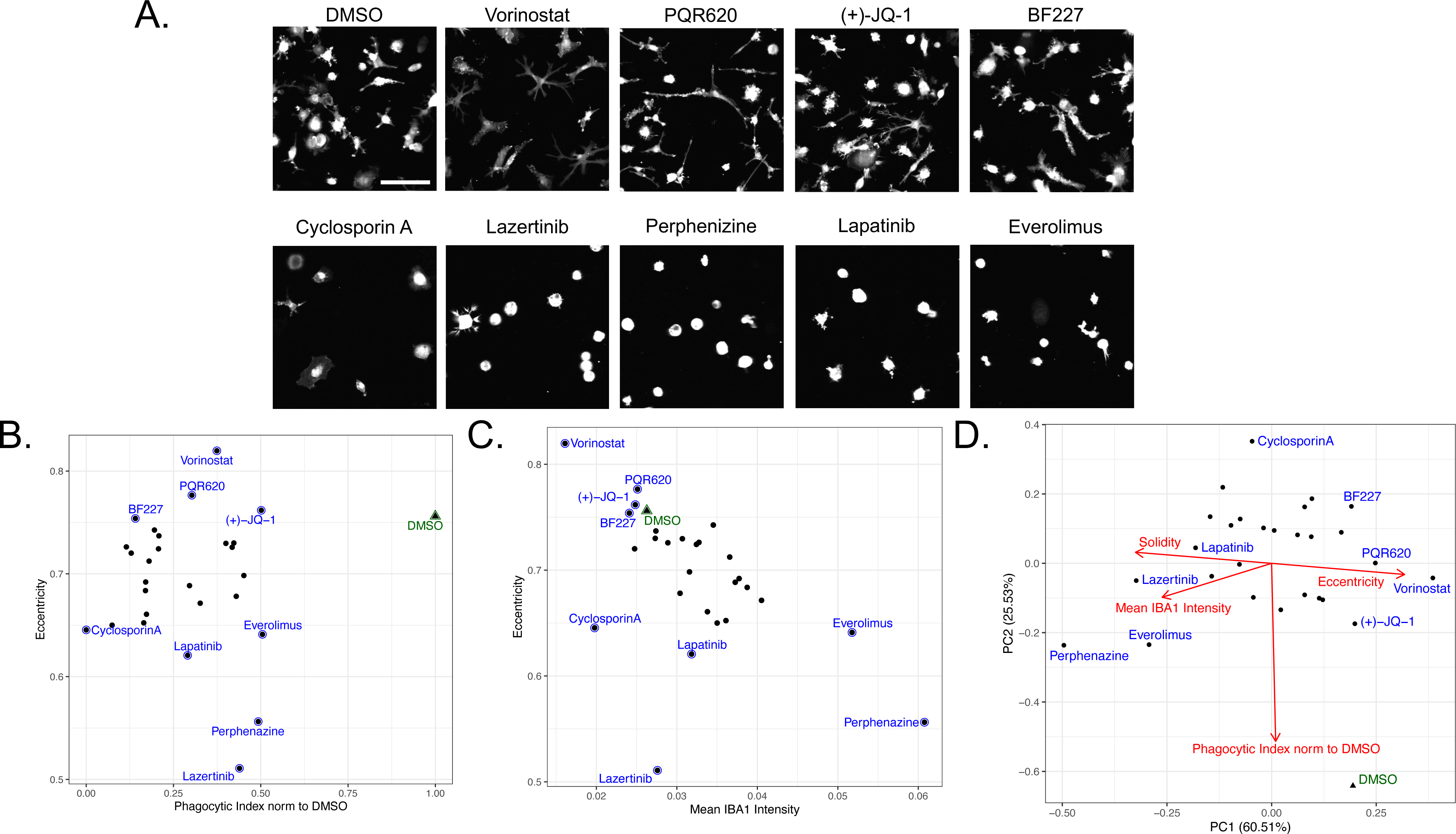
Phenotypic screening identifies compounds that modulate piMGLC morphology. (a) Representative images of DMSO vehicle control and select compound treatments resulting in ramified (top row) or amoeboid (bottom row) morphologies. (b) Phagocytic index versus eccentricity for compounds ≤ 50% DMSO control phagocytic index in secondary screen. (c) IBA1 intensity versus eccentricity for compounds ≤ 50% DMSO control phagocytic index in secondary screen. Each dot in (b) and (c) is mean value of 5-15 image fields with indicated compounds corresponding to images in (a) for representation. (d) Principal Component Analysis (PCA) of functional and morphological features over compounds exhibiting ≤ 50% DMSO control phagocytic index. PC loading vectors in red demonstrate the relationship between features.

Compounds that reduced phagocytosis by 50% or more in the secondary confirmation screen resulted in varied responses in cell morphology (Fig. 4A and Supplemental Data Table 4). For example, though they share low phagocytic indices, vorinostat treatment increased cell ramification as shown by increased eccentricity (Fig. 4B and Supplemental Fig. S1A) and reduced solidity (Supplemental Fig. S1B) compared to the DMSO negative control, whereas perphenazine treatment conversely resulted in exclusively compact amoeboid morphology. Cyclosporin A, the compound resulting in the greatest phagocytic inhibition upon treatment, also decreased eccentricity though not to the same extent. Taken together, these observations suggest the mechanisms by which these compounds modulate phagocytosis of synaptosomes have divergent effects on microglia morphology and, by proxy, activation state.

We also examined IBA1 intensity during the secondary screening as a potential indicator of activation state. While IBA1 is a widely used marker to identify microglia, it is not exclusively tied to activation as it is constitutively expressed in these cells under both resting and activated conditions. However, increased IBA1 levels may indicate more activated microglia when taken in conjunction with other parameters such as phagocytic activity and morphology^38,39^, or additional activation markers.^40^ To this end, we further analyzed the correlation between these parameters in our image-based secondary screen. With a few exceptions, such as lazertinib and cyclosporin A, we observed that compound treatments resulting in higher eccentricity morphology corresponded with lower IBA1 intensity (R=-0.565, p=0.0014) (Fig. 4C). The opposing trend was observed using the related parameter of solidity (R=0.624, p=0.0003) (Supplemental Fig. S2B). The relationship between multiple morphological and phenotypic variables can be summarized using Principal Component Analysis (Fig. 4D). As expected, solidity and eccentricity are highly negatively correlated, as indicated by parallel and opposite principal component loading vectors. Conversely, the Phagocytic Index loading vector appears as nearly perpendicular to the solidity-eccentricity axis, indicating that it is uncorrelated. Mean IBA1 intensity correlates most strongly with solidity, as shown in Fig. 4D.

Taken together, further phenotypic characterization of the 28 compounds prioritized in our secondary confirmation screening shows differential effects on morphology as a proxy for cell state, suggesting different mechanisms resulting in decreased phagocytic activity.

### Transcriptomics identifies divergent pathways affecting phagocytosis and activation state

To investigate potential mechanism of action of the 28 compounds confirmed in the secondary screen, we performed DRUG-seq^41^, a high-throughput multiplexed next-generation RNA sequencing method to measure the effect of 24-hour compound treatment on cellular transcriptional programs in piMGLCs. In all, 16/28 compounds were determined to have measurable effect on transcriptional activity (“RNA-active”, see Methods), with the highest number of differentially expressed genes (DEG) observed in vorinostat-treated wells (3525) (Fig. 5A). We visualized the grouping of replicate wells using Uniform Manifold Approximation and Projection (UMAP), which demonstrates reproducibility of the treatment effects across wells and batches, with tighter grouping of replicates observed for compounds which had a stronger effect on transcription (Fig. 5B). Functional enrichment analysis performed with QIAGEN Ingenuity Pathway Analysis (IPA) shows diverse effects of the compounds (select categories shown in Fig. 6A, full list given in Supplemental Data Table S6.

**Figure 5.**
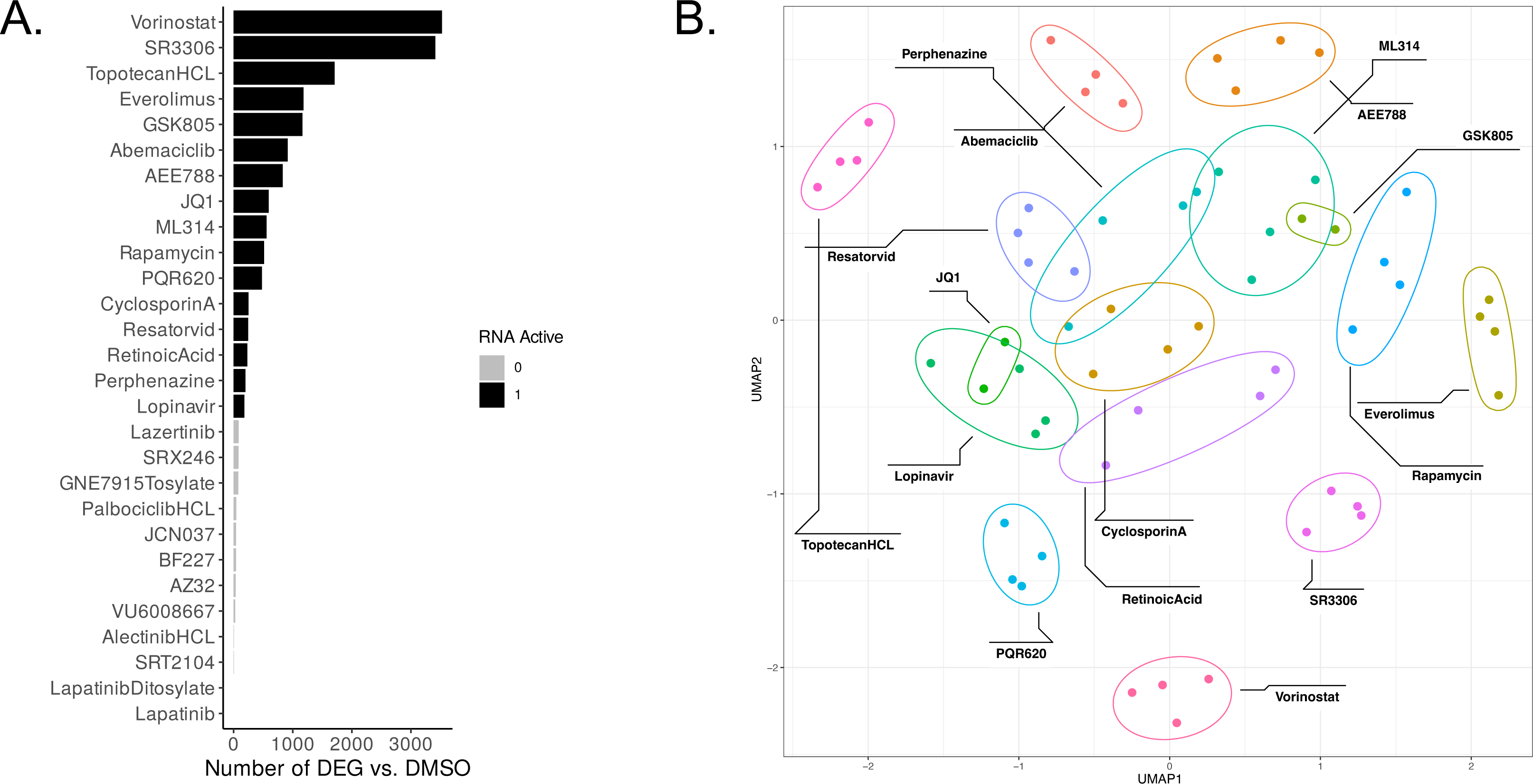
DrugSeq transcriptomic results for secondary screen. (a) Number of differentially expressed genes (DEG) resulting from the comparison of all replicates of a given compound vs. select DMSO replicates (see Methods). Active RNA compounds had more DEG than 95% of the DMSO vs. DMSO comparisons (see Methods). (b) UMAP of active RNA compounds shows grouping of replicates.

**Figure 6.**
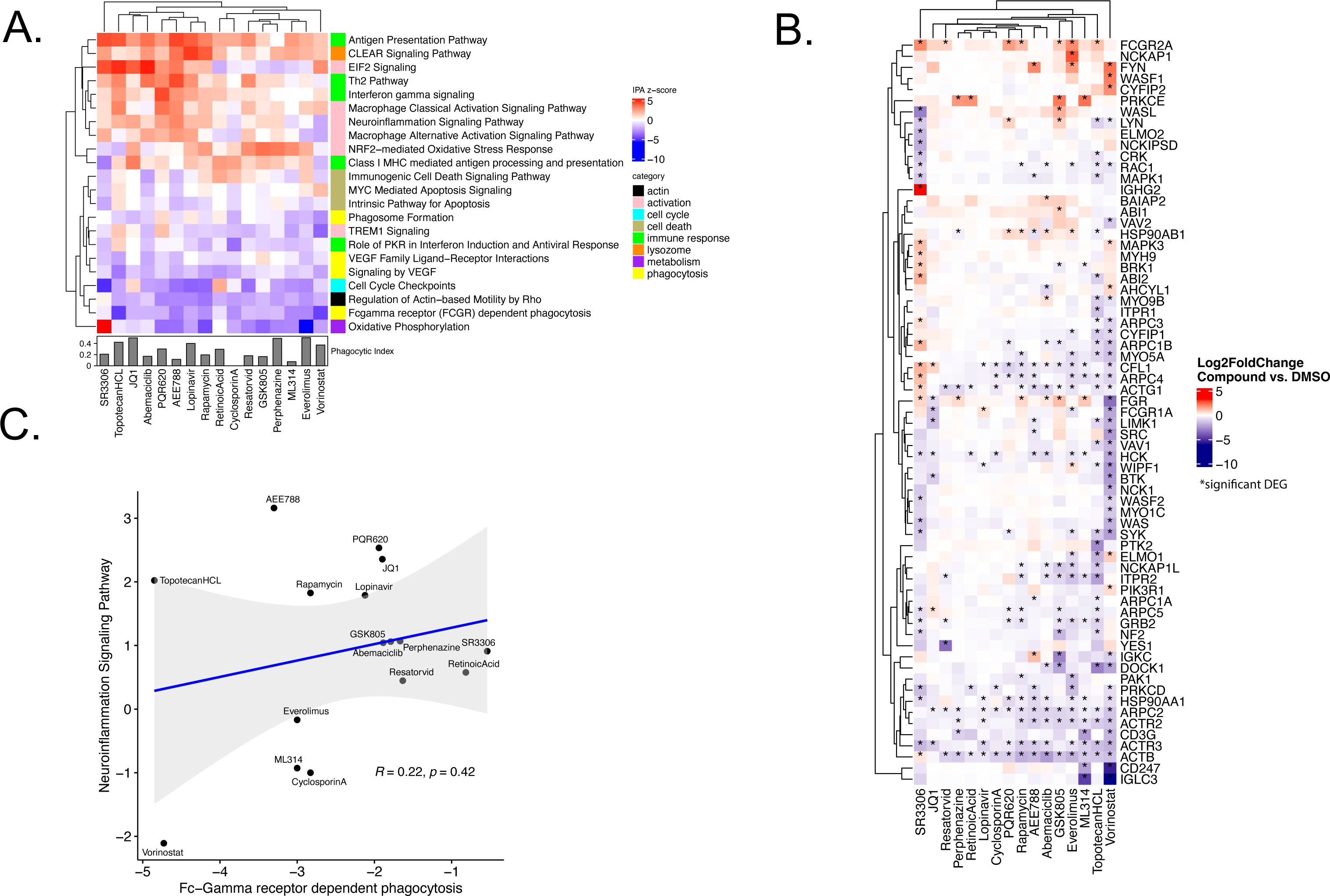
Transcriptomic pathway analysis demonstrates the diversity of compound treatment responses. (a) Activation/Suppression z-score for select IPA Canonical Pathways that are enriched in the DEG of DrugSeq compounds. (b) Relationship between phagocytosis and neuroinflammation for RNA-active compounds exhibiting ≤ 50% DMSO control phagocytic index showing the unique position of Vorinostat. (c) Heatmap of gene differential expression in the FCGR-dependent phagocytosis IPA Pathway by compound. Color represents gene expression level (log_2_ fold change relative to DMSO), *indicated a significant DEG with adjusted p-value < 0.1 and absolute log2 fold-change > 1.

All compounds suppressed gene pathways involved in Fc gamma receptor-dependent phagocytosis, most strongly vorinostat, and most suppressed phagosome formation, with the exception of SR3306, PQR620, lopinavir, topotecan HCL, and retinoic acid. A closer examination of differentially expressed genes in the Fc-gamma receptor (FCGR)-dependent phagocytosis pathway provides insights into the distinct mechanisms by which the compounds reduce phagocytosis (Fig. 6B). Genes involved in the regulation of actin cytoskeleton were most uniformly downregulated, including beta (β)-actin (*ACTB*) and actin-related proteins *ACTR2* and *ACTR3*, as well as heat shock protein *HSP90AA1*.^42^ Cytoplasmic FMR1 interacting protein (CYFIP) family genes *CYFIP1* and *CYFIP2*, involved in actin polymerization, were also impacted, but in divergent ways across compounds. *CYFIP1*, which we recently demonstrated to be involved in microglial phagocytosis, morphology and motility^31^, was downregulated in 3/16, including vorinostat, which strongly upregulated *CYFIP2*. FCGR genes were also differentially impacted across compounds. *FGR* was upregulated in 6/16 and down in 2/16, including vorinostat. *FCGR1A* was downregulated in 3/16 and upregulated in only lopinavir. *FCG2RA* was upregulated in 7/16, most strongly in everolimus, SR3306 and PQR620.

Regulation of actin-based mobility by Rho was also suppressed by all compounds except SR3306, and the pattern of suppression closely mirrored the suppression of phagocytosis. This is consistent with the established role of actin remodeling in phagocytosis.^43^ VEGF signaling pathways were similarly broadly suppressed across compounds. In contrast, cell activation pathways featured prominently among activated pathways, including Classical and Alternative Macrophage Activation, Neuroinflammation, EIF2 Signaling, NRF2-mediated Oxidative Stress Response, and TREM1 Signaling. The degree of activation varied among compounds with the strongest overall activation observed for SR3306, topotecan HCL, JQ1, abemaciclib, PQR620, AEE788 and lopinavir. In a less activating group of compounds including cyclosporin A, resatorvid, GSK805, perphenazine, everolimus and ML314, EIF2 signaling was decreased, while NRF2-mediated oxidative stress response was increased. This pattern suggests a potentially balancing interaction between the autophagy and oxidative stress responses.^44^

Antigen presentation pathways were activated to varied extents across compounds, except for suppression of Class I MHC mediated antigen processing and presentation in SR3306. SR3306 overall showed the most distinct response from the group of RNA-active compounds, with strongly activated Oxidative Phosphorylation and strongly suppressed Cell-cycle checkpoint pathways.

A highly promising candidate therapeutic is one that could maximize suppression of phagocytosis without causing high levels of cell activation, minimizing the risk of off-target effects on microglia. Although there is no strong relationship between the IPA z-scores for FCGR dependent phagocytosis and neuroinflammation across compounds (Fig. 6C), vorinostat was identified as one such compound which strongly suppressed FCGR-dependent phagocytosis without increasing neuroinflammation.

## Discussion

In this functional compound screening of a CNS-penetrant compound library we initially identified 47 compounds that modulate microglia-mediated phagocytosis of synaptosomes in an *in vitro* model of synaptic pruning^14,27,30^, of which 28 demonstrated consistent inhibitory effects in secondary confirmation screening. Notably, while these confirmed hit compounds all inhibit phagocytosis in microglia-like cells, they exhibit variability in additional phenotypic screening, including differential effects on cellular morphology and transcriptomic effects, indicating multiple mechanisms of action rather than reflecting a single common pathway. In fact, these compounds cover a wide range of therapeutic classes each with different mechanisms of action: immunosuppressants, kinase inhibitors, neurotransmitter modulators including antipsychotics and epigenetic modulators. Despite these mechanistic differences, all the compounds diminish microglial phagocytosis, suggesting the potential to reverse microglia-mediated pruning dysregulation implicated in multiple neurological disorders.^45,46^

Specifically, immunosuppressive compounds confirmed in secondary screening including cyclosporin A, resatorvid and rapamycin, which have been shown to affect microglial activation by inhibiting calcineurin^47^, Toll-like receptor 4 (TLR4)^48^ and mechanistic target of rapamycin (mTOR)^49^, respectively, and reducing pro-inflammatory cytokine production and neuroinflammation. Pathway enrichment analysis of transcriptomic data indicated that compound treatment strongly suppressed Fc gamma-receptor mediated phagocytosis while inducing, to diverse extents, cell activation pathways. This diversity enabled us to select compounds with strong phagocytosis reduction and minimal cell activation.

Similarly, we show that epigenetic modulators, such as the histone deacetylase (HDAC) inhibitor vorinostat, greatly reduce synaptosome phagocytosis and hyper-ramify the piMGLCs with concomitant suppression of, e.g. FCGR-dependent phagocytosis, VEGF signaling, regulation of actin motility, and oxidative phosphorylation. Identified kinase inhibitors including lapatinib, lazertinib, palbociclib and dasatinib target cell signaling pathways that may regulate microglial function and synaptic plasticity.

We further show that antipsychotics identified in our screen, including loxapine and perphenazine which exert their primary antipsychotic effect through neuronal dopamine receptor antagonism, also diminish microglial phagocytosis function. Other identified neurotransmitter modulators such as vilazodone affect the levels of serotonin. While less studied than in their neuronal counterparts, microglia do express serotonergic and dopaminergic receptors and respond to serotonin (HT) and dopamine.^50,51^ Notably, loss-of-function in the 5-HTR_2B_ serotonergic receptor, the predominant HT receptor expressed in microglia, reduced presynaptic material phagocytosis in an engineered mouse model.^52^ Our results suggest that these drugs may exert more direct effects on microglia via these pathways, beyond their known neuronal effects.

Whether these effects contribute to antipsychotic or mood stabilizing effects *in vivo* merits further study.

Finally, we identified compounds targeting the mTOR pathway including rapamycin (sirolimus) and its analog, everolimus, as modulating both synaptic phagocytosis and morphology. These compounds have been shown to reduce microglial pro-inflammatory cytokine production and shift polarization towards an anti-inflammatory state^53,54^, which could reduce synaptic engulfment and neuroinflammation in conditions such as AD^55^, schizophrenia^56^ and bipolar disorder.^57^

Several of the identified compounds have been investigated pre-clinically for neurological conditions including psychiatric and neurodegenerative disorders as well as traumatic brain injury (TBI). Examples include SRX246, a potent and highly selective vasopressin 1a receptor antagonist, studied for its potential effects on stress response and mitigation of anxiety and aggression in Huntington’s Disease (HD) patients^58^; SRT2104, a Silent information regulator 1 (SIRT1) activator that promotes cellular survival pathways, reduces oxidative stress, and may slow neurodegenerative progression in disorders including AD and Parkinson’s^59^; and resatorvid (also known as TAK-242), a TLR4 inhibitor, that has been explored in animal models for its ability to reduce neuroinflammation and protect against brain injury by decreasing microglial activation^60^, as well as clinical studies of severe sepsis where it was shown to be safe and tolerable, though ineffective for this indication.^61^

On the other hand, many of our small molecule screening hits have not previously been investigated clinically for CNS indications. For example, the tyrosine kinase inhibitors lapatinib, alectinib and lazertinib have been extensively investigated, but exclusively in the treatment of various cancers.^62^ Another example is the HDAC inhibitor vorinostat, which has a favorable safety and tolerability profile when used in monotherapy or in conjunction with other therapies for the treatment of cutaneous T-cell lymphoma (CTCL).^63^

While some of these drugs, including vorinostat^64^, have shown promising results in animal models by modulating various microglial functions and potentially impacting neurological pathology, they have not been studied in humans in psychiatric and neurodevelopmental disorders. Randomized controlled trials would be needed to establish the clinical utility and safety profiles of these drugs in treating these disorders.

In aggregate, our results identify a range of CNS-penetrant compounds with established human safety profiles. In light of the link between microglial dysregulation and multiuple neurodevelopmental, neurodegenerative and psychiatric disorders,^21,65,66^ these therefore represent high-priority candidates for further study *in vivo*. More broadly, understanding the precise mechanisms by which these drugs modulate microglial function could open new therapeutic avenues for treating a range of brain diseases. Notably, given their potential to shift pruning trajectories, they may find application not solely in controlling symptoms, but potentially in modifying disease course.

## Methods

### Large-scale isolation of PBMCs from leukapheresis

PBMCs were isolated from a half leukapheresis pack (leukopak) sourced from HemaCare Corporation (now part of Charles River Laboratories Cell Solutions) using a standard operating procedure from the AIDS Clinical Trials Group Laboratory Technologist Committee (https://www.hanc.info/content/dam/hanc/documents/laboratory/actg-impaact-laboratory-manual/Leukopak%20PBMC%20Processing%20Standard%20Operating%20Procedure%20v2.0_22Nov2022.pdf). Contents of the leukopak were diluted with Hank’s Balanced Salt Solution without Calcium, Magnesium, phenol red (HBSS, Thermo Fisher #14175095) using at least a 1:2 ratio of leukopak to dilutant. 50ml conical tubes were prepared by adding 15ml Density Gradient Medium (DGM, Stemcell Technologies Lymphoprep 07851) to the bottom of each tube. The full volume of the diluted leukopak was split evenly into the 50ml conical tubes, layering the solution on top of the DGM in each tube. Tubes were centrifuged at 400g for 30 minutes, with centrifuge brake set to “off”. The PBMC fractions between the plasma and DGM were transferred to new tubes, and diluted with HBSS to a final volume of 45ml each. A series of washing steps followed, starting with a centrifugation at 300g for 10 minutes, again with no brake. The supernatants were discarded, and pellets resuspended in 5ml HBSS, before combining into 2 x 50ml conical tubes. This wash was repeated, and cells were consolidated into a single conical tube with a 20ml volume. Cell concentration was quantified using a Countess II automated cell counter (ThermoFisher), and the volume of Cryostor cs10 medium (Sigma-Aldrich C2874) was calculated to aliquot PBMC’s at 25 million cells/ml. The final centrifugation at 300g for 10 minutes was completed and the pellet was resuspended in the Cryostor cs10 medium, before slow freezing in a Mr. Frosty (ThermoFisher #5100-0001) freezing container at -80C and moving to long term liquid Nitrogen storage the following day. The cells were suspended in Cryostor cs10 at room temperature for a maximum of 10 minutes before being added to the freezer. If the time to aliquot this suspension was estimated to be longer, the cell solution was split into multiple batches before the final centrifugation, and aliquots were prepared one batch at a time.

### PBMC-derived induced microglia-like cell (piMGLC) batch culture and assay plate seeding

Frozen and aliquoted peripheral blood mononuclear cells (PBMCs) isolated as described above from a healthy control donated half leukopak (Charles River, Lowell MA / Hemacare, Northridge CA) were quick-thawed in a 37C water bath and immediately transferred into RPMI-1640 (Sigma, #R8758) + 10% heat-inactivated fetal bovine serum (Sigma, #12306C) + 1% of Penicillin/Streptomycin (Life Technologies, cat# 15140-122). Cells were washed by centrifugation at 300 g for 5 min with the brake off and resuspended in fresh media, then counted and plated at a density of approximately 400,000 cells/cm^2^ in a tissue culture treated 6-well plate (Corning, #353046) pre-coated with Geltrex (Gibco, #A1413202) for 1 hour. After incubating for 24 hours, the media was carefully removed and replaced with RPMI-1640 + 1% Penicillin/Streptomycin + 1% Glutamax (Life Technologies, # 35050-061) + 100 ng/ml IL-34 (Biolegend Inc, #577904) + 10 ng/ml GM-CSF (PeproTech, #300-03). After a 10-day incubation period for trans-differentiation, culture media was collected and filtered with a 0.22um Steriflip-GP Filter (EMD Millpore #SCGP00525) and retained. PBMC-derived induced microglia-like cells (piMGLCs) were harvested using Accutase (Sigma, #A6964) for 5 minutes at 37C to detach. They were then centrifuged at 300g for 5 min and counted so they could be resuspended in the proper amount of retained and filtered cell culture media to 50,000 cells/ml, and plated into 96-well tissue culture plates (Corning, #3904) at a density of 30,000 cells/cm^2^ (10,000 cells per well in 200ul) for 4 days before assaying. If the assay plates required a higher volume of cell-culture media than that collected from the batch of 6-well plates, supplementary fresh media of RPMI-1640 with Penicillin/Streptomycin and Glutamax would be used to reach the required seeding volume.

### Compound treatments

24 hours before the start of the assay, the media volume per well was brought to 150ul and piMGLCs were pretreated with the 489 screening library compounds (CNS-penetrant compound library, MedChemExpress #HY-LO28) (Supplemental Data Table S1) at a final concentration of 10μM, with some exceptions at 2μM (Supplemental Data Table S2), by diluting the compounds in basal RPMI-1640 and adding 50ul of one diluted compound per well. For the primary screen a single dilution per compound was used in a single well per plate, with three replicates split across different plates. DMSO (0.1%) was used as a vehicle control and was used in 3 replicate wells on each plate. The secondary screen included piMGLC plates treated with 47 compounds for 24-hours (Supplemental Data Table S4).

### Phagocytosis assays

Phagocytosis assays were performed in 96-well plates (Corning, #3904) containing piMGLCs at a density of 30,000 cells/cm^2^ (10,000 cells per well in 200ul). Human synaptosomes were thawed at room temperature and an equal volume of 0.1M sodium bicarbonate, pH 9 was added to the synaptosomes. pHrodo-Red (Invitrogen, #P36600, 6.67ug/ul) was added at a protein ug ratio of 1:2 (dye:synaptosome). The labeling reaction was incubated at room temperature for 1 hour in the dark. Synaptosomes were washed by adding at least an equal volume of PBS, pH 7.4 to the tube and pelleting at 12,000 rpm for 15 min. The wash solution was discarded and the labeled synaptosomes were resuspended in a volume of basal RPMI-1640 to a final concentration of 0.15 μg synaptosomes per μl. Labeled synaptosomes were sonicated in a Branson 1800 (Emerson, #M1800) at 40 kHz for 1 hour. During this time, some wells of microglia were treated with cytochalasin-D (Sigma, #C2618) to a final concentration of 2 μM as a control treatment to inhibit phagocytosis. Following sonication, the synaptosomes were added to wells at a final concentration of 3μg/well. Due to the scale of the primary screen, synaptosomes were added to plate wells using a Multidrop Combi reagent dispenser for this experiment (Thermo Scientific, 5840340). Synaptosome were added to wells using a multichannel pipette for experiments following the primary screen. Phagocytosis assays were ended after 3 hours by fixation with 4% paraformaldehyde (Electron Microscopy Sciences, #15713S).

### High Content Image Analysis

Fifteen randomly spaced confocal microscopy images per well were analyzed using CellProfiler (Version 4.2.1).^32^ Specific pipeline parameters used are detailed in Supplemental Data Table S7. Briefly, illumination correction was used to decrease background signal in the images, nuclei and piMGLC structures were segmented and masked to each other, synaptosomes were segmented and masked to the cytoplasm, and all descriptive calculations such as signal intensity and object size for each of the identified objects were exported. The pipeline was fine-tuned to ensure optimal object segmentation accuracy, and steps were added to the downstream analysis to handle any object segmentation errors that did occur. The same pipeline was used for the analysis of all experiments, and only the lower bounds of the automated Otsu thresholding for synaptosome segmentation were adjusted between experiments since each may have a different background fluorescent signal intensity. To set the optimal lower bound on the thresholding and prevent background fluorescence from being included in synaptosome segmentation, 2uM cytochalasin D treated wells (phagocytosis inhibitor used as positive control) were included in each plate to have a low signal well to calibrate the thresholding bounds.

Field level filters were applied for identified cell number and PI, next fields were aggregated to wells and similar well-level filters were applied. Data exported from CellProfiler were loaded into an R notebook for cleaning. Initially, cell number was used to identify toxic treatments. Further, we identified potential CellProfiler segmentation errors. Next, a filter was applied to retain only images with between 4 and 80 cells, thus omitting the extremely dense or sparse fields that likely result from CellProfiler over-segmentation or treatment toxicity, respectively. The next filter removed image fields where the phagocytic index (synaptosome area divided by cell count) was greater than the mean plus three times the standard deviation (3SD), which would be a result of inaccurate synaptosome segmentation.

The secondary confirmatory compound screen used the image derived data at this more sensitive image field granularity, with a minimum of 5 image fields remaining after the cleaning steps noted above. The primary screen used well-level quantification to reduce noise in the larger scale experiment. To do this the well-level phagocytic index was calculated as the sum of synaptosome area divided by the sum of cells in all images per well. Similar cleaning filters were applied to the well-level data. First, the same max phagocytic index threshold of mean plus 3SD was used. Along with this, wells with low per-well cell counts were omitted using a minimum count threshold of 10% of the mean of cell counts in DMSO treated wells. Compounds with fewer than three remaining replicate wells after applying these filters were omitted from further analysis.

### Multiplexed library preparation and RNA sequencing

96-well multiplexed libraries (DRUG-seq ^41^) were prepared using the MERCURIUS^™^ DRUG-seq kit (Alithea Genomics, 10841) following manufacturer instructions. Briefly, cells were washed using 100µL per well of Dulbecco’s Phosphate Buffered Saline (Gibco, 14190-144). Cells were then lysed for 10-15 minutes in 20µL per well of prepared and chilled 1x Cell Lysis Buffer at 4°C. Lysate was carefully removed from the cell culture plate and transferred to a 96-well PCR plate, then spun at 300xg for 5 min in a centrifuge pre-chilled to 4°C. Up to 20µL of supernatant was then removed to a clean 96-well PCR plate for storage at -80°C for up to one month.

For reverse transcription, stored lysate was thawed at 4°C, then 10µL of lysate from each well was transferred to a 96-well plate containing dry, well position specific barcoded oligo-dT primers (Alithea Genomics, 10513). Upon resuspension of the oligo-dT primers, 10µL of prepared RT Master Mix was added to each well and reverse transcription was carried according to provided instructions (incubation for 30min at 50°C, 10min at 85°C, then cooling to 4°C). Resulting first-strand cDNA was collected and pooled, mixed with 7x volume of DNA binding buffer (Zymo, D4003-1-L), and purified on a Zymo-Spin IC column (Zymo, C1004-50). Purified first-strand cDNA was eluted in 20µL nuclease-free water. Excess oligo-dT was digested from 17µL of eluate by adding Exonuclease I Enzyme and Buffer and incubating for 30min at 37°C, 20 min at 80°C, then cooling to 4°C. Second-strand synthesis was carried out immediately after by adding 7µL of prepared Second Strand Synthesis reaction mix and incubating for 20 min at 37°C, 30 min at 65°C, and then cooling to 4°C.

Resulting second-strand cDNA was purified using CleanNGS DNA & RNA Clean-Up Magnetic Beads (Bulldog Bio, CNGS005) at a ratio of 0.6x bead slurry to cDNA. Bead purification was carried out using a 0.2 mL PCR Strip Magnetic Separator (Permagen, MSR812). Upon purification, cDNA was measured using Qubit 1X dsDNA HS Assay kit (Invitrogen, Q33231) and 50-60ng of cDNA was tagmented in a 20µL reaction with 4µL of provided Tagmentation Enzyme and 4µL of Tagmentation Buffer. After incubating at 55°C for 7 minutes, the tagmented library was brought to 50µL and purified with CleanNGS beads at a ratio of 0.6x bead slurry to cDNA. Finally, the tagmented library was pre-amplified with provided UDI adapters included in kit for 10 cycles (one cycle composed of 10 seconds at 98°C, 30 seconds at 63°C, and 1 min at 72°C) total. In preparation for sequencing, this pre-amplified library was purified twice with CleanNGS beads at a ratio of 0.7x bead slurry to cDNA and eluted in 20uL.

Quality control and measurement of library concentration of finished libraries was performed with Agilent 5300 Fragment Analyzer System using the HS NGS Fragment Kit (Agilent, DNF-474-0500), for quality control and measurement of library concentration. Library was loaded onto one lane of a NovaSeq X Plus 1.5B read flow cell and sequenced on the NovaSeq X Plus instrument. Sequencing cycle structure was as recommended by manufacturer (Read 1: 28 cycles; i7 read: 8 cycles; i5 read: 8 cycles; Read 2: 90 cycles). Samples were split over two different sequencing runs, with at least one replicate of each studied compound on each run. Sequencing resulted in 615 million read pairs for run one and 1016 million read pairs for run two.

### DRUG-seq data analysis

Sample demultiplexing and read alignment was done following the data analysis pipeline steps in MERCURIUS DRUG-Seq kit documentation. Briefly, raw reads were aligned to reference genome hg38 using STAR v. 2.7.11b ^67^ using “solo” mode. Unique molecular identifier count matrices were generated in this step by supplying the list of sequencing barcodes to option – soloCBwhitelist (Supplemental Data Table S8). The resulting demultiplexed count matrix contained on average 3 million ± 1 million reads per sample and was used for downstream analysis.

Differential Expression analysis was done using a modified version of the DRUG-seq analysis pipeline ^68^ using DESeq2 ^69^ to normalize the count matrix, fit a negative binomial regression model with design = ∼ batch + compound, and perform differential expression analysis using the Wald test. Genes were considered significantly differentially expressed with adjusted p-value < 0.1 and absolute log2 fold-change > 1, using each compound and DMSO as comparison groups. Briefly, using all DMSO wells, we determine the “best” DMSO wells by performing 500 random DMSO vs. DMSO comparison, and selecting the two DMSO replicates from each group that contributed to the fewest differentially expressed genes (DEG). Next, we determine the null distribution of DEG by comparing 500 randomly sampled sets of DMSO replicates to our chosen 4 “best” DMSO wells. Finally, we compare all replicates of each treatment compound vs. the 4 best DMSO wells. Treatment compounds were considered “RNA active” if the number of DEG resulting from the comparison with DMSO was greater than the 95% of DMSO vs. DMSO comparisons, which was determined to be 179 DEG. Uniform Manifold Approximation and Projection (UMAP) dimension reduction was used to visualize overall similarity of transcriptional profile of all transcriptionally active treatment wells. Functional enrichment analysis for all sets of treatment compound vs. DMSO comparisons DEGs was performed using QIAGEN Ingenuity Pathway Analysis v. 24.0.1 ^70^ canonical pathway analysis.

## Data Availability

Gene expression data will be made available for download from the NCBI Gene Expression Omnibus (https://www.ncbi.nlm.nih.gov/geo) upon publication. Additional data that support the findings of this study will be available from the corresponding authors upon reasonable request.

Additional assay details available in Supplemental Material and Methods

## Supporting information

Description of Additional Supplemental Data Tables

Supplemental Data Table S1 Full Library Information

Supplemental Data Table S2 Other Drug Treatment Info

Supplemental Data Table S3 Primary Screen PI and Z-score Results

Supplemental Data Table S4 Secondary Screen Results

Supplemental Data Table S5 Secondary Screen Compound Clinical Status

Supplemental Data Table S6 IPA Results Full List

Supplemental Data Table S7 CellProfiler Pipeline Settings

Supplemental Data Table S8 Drug-Seq Treatments and Barcodes

## Acknowledgements

This work was supported by R01MH120227 (Perlis) and R01MH131687 (Sheridan and Perlis). The next generation sequencing was performed in Tufts University Core Facility Genomic Core, which received funding support from NIH (S10 OD 032203) for its Illumina NovaSeq sequencer.

## Author Contributions

L.T.M. and T.F. generated, purified, and characterized large scale synaptosome preparations. L.T.M, H.Y. and J.M.T. optimized the batch differentiation and characterization of piMGLCs, and scaling phagocytosis assay to 96-well format. L.T.M. performed the phagocytosis drug screens, morphological assays, developed CellProfiler pipelines and data analysis. R.E.B. performed transcriptomic, morphology, principal component analyses and interpretation of data. J.J.B. performed sample and multiplexed Drug-seq library preparation for sequencing. S.D.S., R.E.B., L.T.M and R.H.P wrote the manuscript. S.D.S and R.H.P conceived and directed the project. All authors have read and approved the manuscript for publication.

## Materials & Correspondence

Correspondence and material requests should be addressed to R.H.P or S.D.S

