## Supplemental Materials and Methods for "Identifying brain-penetrant small molecule modulators of human microglia using a cellular model of synaptic pruning"

Running Title: Small molecule modulators of human microglia

### Supplemental Materials and Methods

### Supplemental Figures:

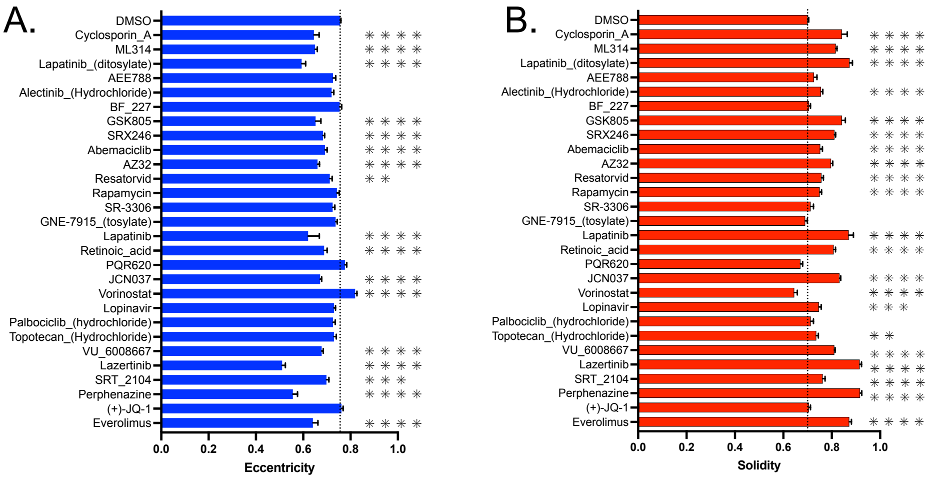


**Figure S1**. Bars represent the means of image-level piMGLCs (a) eccentricity and (b) solidity values for the 28 compounds confirmed to decrease phagocytosis by 50% or more in the secondary screen. The vertical dotted line represents the DMSO mean value, asterisks signify the adjusted p-values of the dunnett's multiple comparison test, which tests each compound's value to the DMSO control. One to four asterisks correlate to an adjusted p-value of <0.05, <0.01, <0.001, and <0.0001, respectively. Error bars indicate SEM.

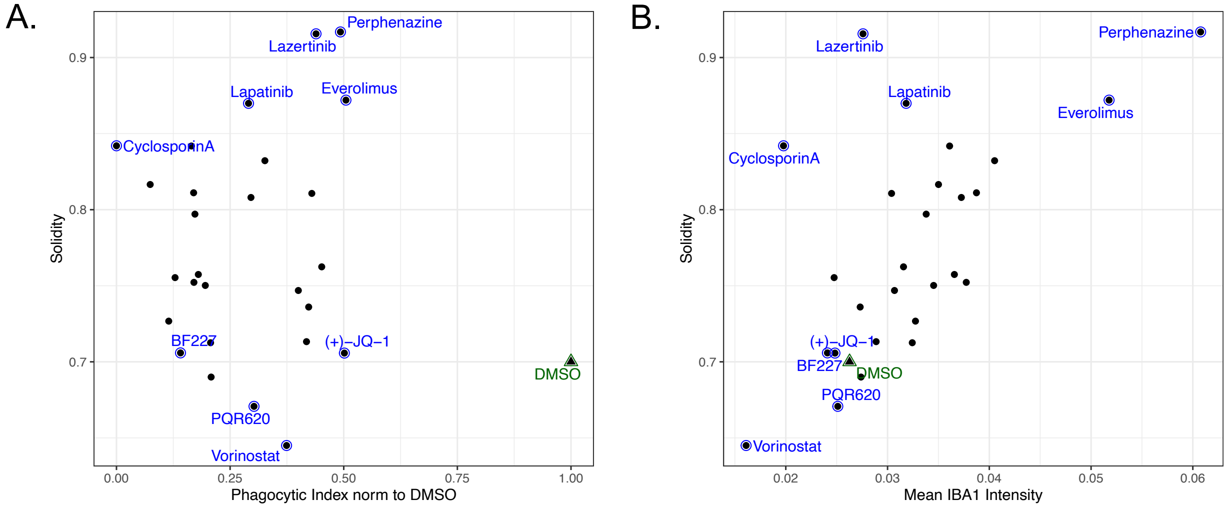


**Figure S2**. Phenotypic screening identifies compounds that modulate piMGLC morphology. (a) Phagocytic index versus solidity for compounds ≤ 50% DMSO control phagocytic index in secondary screen. (b) IBA1 intensity versus eccentricity for compounds ≤ 50% DMSO control phagocytic index in secondary screen. Each dot in (a) and (b) is mean value of 15 image fields with indicated compounds corresponding to images in Figure 4A for representation.

### Supplemental Methods:

#### Large scale generation of synaptosomes from iPSC-derived neural cultures for use in models of synaptic pruning screening assays.

iPSCs were reprogrammed from fibroblasts and used to derive neural progenitor cells (NPCs), which were differentiated into neural cultures, as previously described.[1-4] Human iPSC-derived neural progenitor cells (NPC's) cultured in neural expansion media (50% Advanced DMEM/F12, 50% Neurobasal, and neural induction supplement – Gibco: 21103-049, 12634-010, A16477-01) were expanded and seeded (50 million per flask) into T1000 flasks (Millipore, #PFHYS1008) for large-scale neuronal differentiation. NPCs were differentiated in neuronal differentiation medium (Neurobasal media (Gibco # 21103049) supplemented with 1× each (N2 supplement (Stemcell Technologies SCT # 7156), B27 supplement without Vitamin A (Gibco # 12587010), non-essential amino acids (NEAA Gibco # 11140050), penn/strep, 1 µM ascorbic acid, 10 ng/mL BDNF and GDNF (Peprotech), and 1 µg/mL mouse laminin (Sigma # L2020) replacing media weekly for 8-weeks. 24 hours before isolation, media was replaced with human Astrocyte-conditioned medium (ScienceCell, #1811). To start the isolation, media was aspirated and replaced with 1X gradient buffer (0.32M sucrose, 0.75mM $NaHCO_3$, 5mM Tris in Milli-Q water) + HALT protease and phosphatase inhibitor cocktail (Thermo Scientific, #78440), before shaking the flasks to detach the neural cultures from the surface layers of the flask. Cell suspensions were transferred to a dounce homogenizer (Bellco Glass, #1984-10015) and homogenized using 12 up and down strokes per 10ml using the tight plunger. Homogenate was split between 15ml conical tubes and centrifuged at 700g, 4°C, for 10 minutes. Supernatants were saved and the pellets were resuspended in 1X gradient buffer to repeat the previous step once more. Supernatants were combined and transferred to high-speed centrifuge tubes (Beckman Coulter, #344058) to spin at 15000g, 4°C, for 15 minutes. Sucrose gradients were prepared in ultracentrifuge tubes by layering 0.85M sucrose on top of 1.28M sucrose. After centrifugation, the pellets were resuspended in 1X gradient buffer and carefully added to the ultracentrifuge tubes as the top layer of the sucrose gradients. Gradients were centrifuged at 26500 rpm in an ultracentrifuge (Beckman Coulter optima L-90K) using a pre-chilled rotor and swinging cups (SW 32 Ti) for 2 hours at 4°C, with no brake. To collect the synaptosome fraction between the 1.28M and 0.85M sucrose layers, a 5ml syringe with 16G needle was used to pierce the wall of the tube at the level of the synaptosome band to aspirate the fraction, taking as little surrounding gradient buffer as possible. The collected fractions were diluted (at least 1:5) with 1mM $NaHCO_3$ Milli-Q water and centrifuged a final time at 20000g, 4°C, for 20 minutes. Final pellets were resuspended in 1X gradient buffer + 1mg/ml BSA, and aliquoted into cryovials at 100µl/tube. Protein concentration of isolated synaptosomes was quantified using a Pierce BCA protein assay kit (Thermo Scientific, #23225).

**Immunofluorescence**

Live cells were fixed with 4% paraformaldehyde for 15 min at room temperature. Cells in a 96-well format were washed with 100 ul of Wash Buffer, PBS + 0.5% FBS, three times. Then 100 ul of Block and Permeabilization Buffer, PBS + 0.5% FBS + 0.3% Triton-X, were added per well. Wells were incubated at room temperature for 1 hr, then washed three times with Wash Buffer. Primary antibodies were diluted in Antibody buffer, PBS + 0.5% FBS + 0.1% Triton-X, added to the wells and incubated for 1 hr at room temperature or overnight at 4 C. Wells were washed three times with Wash Buffer. Secondary antibodies were diluted in Antibody Buffer and added to wells and incubated for 45 min at 4 C. After a final three washes, the wells were imaged with an IN Cell Analyzer 6000 automated confocal microscope (Cytiva) at 20X.

| Antibody | Species | Vendor (cat#) | Dilution |
|---|---|---|---|
| IBA1 | Chicken monoclonal | Synaptic Systems (234009) | 1:1000 |
| PU.1 | Rabbit monoclonal | Abcam (ab183327) | 1:1000 |
| CX3CR1 | Mouse polyclonal | Abnova (H00001524-B01P) | 1:100 |
| P2RY12 | Rabbit polyclonal | Alamone (APR-020) | 1:100 |
| Chicken secondary | Goat-Alexafluor647 | Invitrogen (A21449) | 1:500 |
| Rabbit secondary | Donkey-Alexafluor488 | Invitrogen (A21206) | 1:500 |
| Mouse secondary | Donkey-Alexafluor488 | Invitrogen (A21202) | 1:500 |

**Morphometric Image Analysis**

Measurements used to characterize cell morphology included eccentricity and solidity. Eccentricity is calculated using an ellipse encompassing the cell and taking the ratio of the distance between ellipse foci and the length of its major axis. This results in a value between 0-1 where long, flatter cells are close to 1, and circular cells are closer to 0.5 (cell morphology never approaches a perfect circle to have a value close to 0).

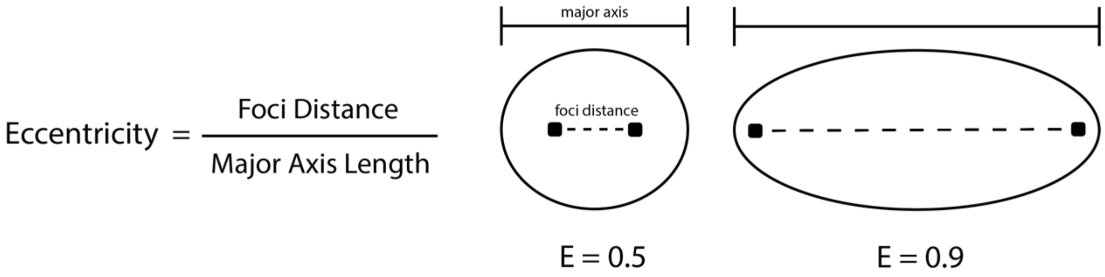


Solidity is calculated using the object's convex hull, the smallest possible convex shape that contains the entire object. Solidity is the ratio of the cell's area to the area of its convex hull, which means more circular cells have a solidity close to 1, while ramified cells with many branches result in solidity values closer to 0.

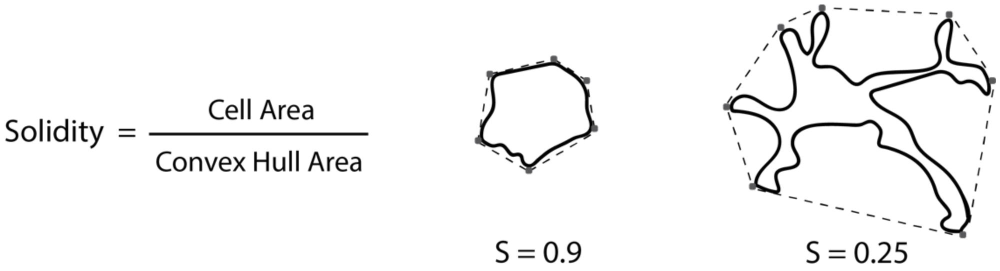
