## Supplementary material for "Identifying brain-penetrant small molecule modulators of human microglia using a cellular model of synaptic pruning": Description of Additional Supplemental Data Tables

**Supplemental Data Table 1**. CNS-penetrant small molecule library of 489 compounds used in primary screen related to Figure 3A.

**Supplemental Data Table 2**. CNS-penetrant small molecule library stocks and final dilution information related to Figs. 3A and 3B and transcriptomic analyses.

**Supplemental Data Table 3**. Normalized mean phagocytic index and Z-score (compared to DMSO control) for primary screen as related to Fig. 3A.

**Supplemental Data Table 4**. Secondary confirmation screen results including phagocytic index (normalized to DMSO control) and measured morphometric parameters: mean eccentricity, solidity and IBA1 intensity related to Figs. 3B, 4B, 4C, 4D, S1 and S2.

**Supplemental Data Table 5**. Clinical phase status of confirmed compounds resulting in <50% phagocytic index reduction related to Fig. 3B.

**Supplemental Data Table 6**. Full list of Ingenuity Pathway Analysis (IPA) implicated pathways related to Fig. 6A.

**Supplemental Data Table 7**. CellProfiler settings used as related to Figs 3 and 4, Figs. S1 and S2 and Methods.

**Supplemental Data Table 8**. Multiplexed Drug-seq barcodes used for transcriptomic analysis related to Figs. 5 and 6 and Methods.
