## Supplemental Data Table S2 Other Drug Treatment Info for "Identifying brain-penetrant small molecule modulators of human microglia using a cellular model of synaptic pruning"

Table S2. Other Drug Treatment Information

All compounds dissolved in DMSO and used at 10uM except for the following:

| Compound Name | Solvent | Concentration |
| --- | --- | --- |
| Primary Screen | | |
| Hoechst 34580 (tetrahydrochloride) | DMSO | 2uM |
| S 38093 | DMSO | 2uM |
| Gatifloxacin | DMSO | 2uM |
| Droxidopa | DMSO | 2uM |
| Itraconazole | DMSO | 2uM |
| SRT 2104 | DMSO | 2uM |
| Tenofovir | DMSO | 2uM |
| Baclofen | DMSO | 2uM |
| Abemaciclib | DMSO | 2uM |
| Palbociclib (hydrochloride) | Water | 10uM |
| Fosphenytoin (disodium) | Water | 10uM |
| Gabapentin | Water | 10uM |
| Secondary Screen | | |
| SRT_2104 | DMSO | 2uM |
| Abemaciclib | DMSO | 2uM |
| Palbociclib_(hydrochloride) | Water | 10uM |
| Drug-Seq | | |
| Abemaciclib | DMSO | 2uM |
